# CDKL5 deficiency alters cell subtype-specific molecular programmes and intercellular signalling in the mouse hippocampus

**DOI:** 10.64898/2026.10.22.760890

**Authors:** Javier Villegas-Salmerón, Samuel Egan, Albert Sanfeliu, Jordan Higgins, Mona Heiland, Bilal El-Mansoury, Niamh M.C. Connolly, David C. Henshall, Omar Mamad

## Abstract

CDKL5 deficiency disorder (CDD) is a severe neurodevelopmental condition characterised by epilepsy, intellectual disability, and motor dysfunction. Loss of CDKL5 leads to altered synapse development, excitatory-inhibitory imbalances and glial changes, but the underlying molecular mechanisms remain incompletely understood, particularly at the level of individual brain cell subtypes. Here, we performed single-nucleus RNA sequencing (snRNA-seq) on the hippocampus of 12 weeks old heterozygous female *Cdkl5* exon 6 deletion (*Cdkl5^tm1.1Joez/J^*) and wild-type mice to characterise cell-subtype-specific transcriptional alterations associated with CDKL5 deficiency.

Analysis of 11,791 high-quality nuclei revealed cell-subtype-specific transcriptional changes across neuronal and non-neuronal populations. Differential expression was most prominent in intratelencephalic and extratelencephalic (IT–ET) glutamatergic neurons, which showed altered expression of genes involved in neuronal signalling and activity-dependent transcription, and in dentate gyrus (DG) neurons, which exhibited coordinated upregulation of multiple ribosomal protein genes. Gene set enrichment analysis revealed broader molecular alterations, with neuronal populations showing recurrent changes in pathways associated with synaptic organisation, neuronal development, axonal and dendritic processes, and neurotransmission. Altered gene expression was also evident among non-neuronal populations, although these displayed fewer changes. CellChat analysis inferred changes in predicted intercellular communication across excitatory and inhibitory neuronal populations, particularly *Sema3c* signalling, suggesting reorganization of hippocampal neuronal subtypes.

These findings provide a single-nucleus transcriptional atlas of the hippocampus in a CDD mouse model. In our dataset, CDKL5 deficiency was associated with cell-subtype-specific alterations in gene expression, including enrichment of gene pathways related to cellular metabolism, neuronal development and synaptic organisation, alongside alteration in predicted intercellular signalling. This dataset provides a resource for investigating the cellular and molecular mechanisms underlying CDD and for prioritising candidate pathways for future therapeutic investigation.

## Introduction

CDKL5 deficiency disorder (CDD) is a severe neurodevelopmental condition characterised by early-onset drug-resistant epilepsy, profound intellectual disability, and motor dysfunction [1–4]. The *CDKL5* gene encodes a serine/threonine kinase with important roles in neurodevelopment and neuronal function, including regulation of neurite outgrowth, dendritic spine development, cytoskeletal and microtubule dynamics, synaptic vesicle function, and transcriptional and epigenetic processes [5,6]. Although our understanding of CDKL5’s functions has grown considerably, its cell-subtype-specific roles and the molecular consequences of its deficiency across distinct neuronal and non-neuronal populations remain incompletely understood [4,7]. Despite the substantial clinical burden of CDD, therapeutic options remain limited, with only one FDA-approved treatment specifically indicated for seizures associated with CDD [8]. This highlights the need for a more detailed understanding of the molecular mechanisms underlying the disorder, particularly the cell subtype-specific consequences of CDKL5 deficiency, to identify potential targets for therapeutic intervention [5].

CDKL5-deficient mouse models recapitulate key features of CDD and provide an important platform for investigating CDD pathophysiology and evaluating novel therapeutics [9]. Bulk transcriptomic and proteomic approaches have previously been used to characterise molecular alterations associated with CDD in mouse brain and patient-derived neuronal models[10–12]. This has revealed dysregulation of pathways involved in neuronal development, cytoskeletal organisation, extracellular matrix interactions and synaptic signalling, while studies using patient-derived neuronal models have identified alterations in neuronal and synaptic regulatory pathways [10,11].

Single-cell and single-nucleus transcriptomic approaches have increasingly revealed cell subtype-specific molecular alterations in experimental epilepsy [13,14], and neurodevelopmental disorders associated with epilepsy, including Rett syndrome [15,16] and HNRNPU-related developmental and epileptic encephalopathy [17]. These have revealed molecular alterations that were obscured by earlier bulk tissue analyses, highlighting the value of single cell approaches to resolve the transcriptional basis of neurodevelopmental disease mechanisms.

Here, we present the first single-nucleus transcriptional atlas of the hippocampus in a mouse model of CDD, using heterozygous female mice carrying a deletion of exon 6 of *Cdkl5*. By resolving transcriptional alterations across individual hippocampal cell populations, this study provides insight into the cell subtype-specific molecular consequences of CDKL5 deficiency and identifies candidate molecular processes for future functional investigation.

## Materials and Methods

### Mouse model of CDKL5 deficiency

All procedures involving animals were performed in compliance with EU Directive 2010/63/EU on the protection of animals used for scientific purposes. Mice were housed under controlled conditions including a room temperature of 20°C–25°C, and humidity (40%–60%), under a 12-h dark/light cycle with *ad libitum* access to water and food.

The mouse model of CDKL5 deficiency disorder used was the B6.129(FVB)-Cdkl5*^tm1.1Joez^*/J (Jackson Laboratory, strain #021967) [18]. The breeding scheme was performed as previously described[19].

### Isolation of nuclei from mouse hippocampi

Two wild-type (WT) and two heterozygous (HET) female mice were anaesthetised at 12 weeks of age and transcardially perfused with PBS prior to brain extraction. Hippocampi were subsequently dissected under sterile conditions. Frozen tissue samples were transferred into 2 mL of Nuclei Extraction Buffer (Miltenyi Biotec, 130-128-024) containing 0.2 U/μL RNase inhibitor (Enzymatics, Y9240L), placed in gentleMACS C Tubes (Miltenyi Biotec, 130-093-23), and dissociated using the gentleMACS Octo Dissociator (Miltenyi Biotec, 130-096-427) with the pre-installed “Nuclei 4C” program. Lysates were filtered through 100 μm MACS SmartStrainers (Miltenyi Biotec, 130-098-463) and centrifuged at 300 × g for 5 min at 4 °C. Pellets were resuspended in 1 mL of Resuspension Buffer (0.5% bovine serum albumin in phosphate-buffered saline with 0.2 U/μL RNase inhibitor) and mixed with 300 μL of Debris Removal Solution (Miltenyi Biotec, 130-109-398) in 2 mL microcentrifuge tubes. Each sample was overlaid with 700 μL of Resuspension Buffer and centrifuged at 3000 × g for 10 min at 4 °C in a swinging-bucket rotor. The upper (debris-containing) layers were carefully aspirated and discarded.

Samples were subsequently washed twice by filling tubes with Resuspension Buffer, inverting three times, centrifuging at 300 × g for 5 min at 4 °C, and removing the supernatant. Nuclei pellets were resuspended in 200 μL of Resuspension Buffer and filtered through 40 μm Flowmi Cell Strainers (SP Bel-Art, H13680-0040). Nuclei concentration and viability were assessed by Trypan Blue staining and manual counting using a hemacytometer.

### Cell barcoding and library preparation

Single-nucleus RNA sequencing (snRNA-seq) libraries were prepared using the Chromium Next GEM Single Cell 3’ Reagent Kit v3.1 (Dual Index) (10x Genomics) on the Chromium Controller, following the manufacturer’s protocol and targeting recovery of ∼3,000 nuclei per sample. Libraries were quantified using the Qubit 4 Fluorometer (Invitrogen, Q33238) with the Qubit 1X dsDNA HS Assay Kit (Invitrogen, Q33231). Quality control was performed with the Agilent 2100 Bioanalyzer using the High Sensitivity DNA Kit (Agilent, 5067-4626).

### Sequencing and primary data processing

Libraries were sequenced on an Illumina NovaSeq platform with a target depth of 100 million paired-end reads (150 bp) per sample. FASTQ files were aligned to the mouse reference genome (GRCm39) and gene expression quantification was performed using Cell Ranger [20] count with the --include-introns=true option enabled.

### Data pre-processing and quality control

All analyses were conducted in R (v4.4.2). Gene expression count matrices were merged using Seurat v5.3.0 [21]. Nuclei with fewer than 500 detected genes or >1% mitochondrial transcript content were excluded. Ambient RNA contamination was corrected using SoupX [22], and doublets were identified and removed using DoubletFinder[23]. Following quality control, 11,791 high-quality single-nucleus transcriptomes were retained for downstream analysis.

### Dimensionality reduction and nuclei labelling

Gene expression data were normalized using Seurat’s NormalizeData() function, and highly variable features were identified using FindVariableFeatures() with default parameters. Expression data were scaled and subjected to principal component analysis (PCA). The number of principal components retained for downstream analysis was selected based on inspection of an elbow plot, with the first 10 principal components used for dimensionality reduction and clustering. Uniform Manifold Approximation and Projection (UMAP) was then performed to visualize nuclei in two-dimensional space and assess sample-level distribution and potential batch effects.

Cell subtype annotation was performed with the MapMyCells software (RRID: <u>SCR_024672</u>) [24], using the 10x Whole Mouse Brain as reference with hierarchical mapping for label transfer. This confirmed the presence of canonical hippocampal cell subtypes, and 21 subtypes were retained for analysis.

### Cell subtype-specific effect of a CDKL5 mutation

To examine the effects of the CDKL5 loss-of-function mutation on gene expression at the cell subtype level, we performed differential expression analysis using two complementary approaches. First, differentially expressed genes (DEGs) were identified using Seurat’s FindMarkers() function with MAST [25] (test.use = "MAST"). Genes with Benjamini–Hochberg-adjusted *p*-values < 0.05 were considered statistically significant.

In parallel, we performed a pseudobulk differential expression analysis. For each cell subtype, raw counts were aggregated by biological replicate using Seurat’s AggregateExpression() function. Differential expression between HET and WILD samples was then assessed using DESeq2[26], with the contrast specified as c("group", "HET", "WILD"). Genes with Benjamini–Hochberg-adjusted *p*-values < 0.05 were considered statistically significant. Principal component analysis (PCA) was performed separately for each cell subtype using variance-stabilized pseudobulk expression data generated with DESeq2 [26]. Genes with a mean normalized count <1 or zero variance were excluded prior to PCA, which was performed using the prcomp() function in R.

### Gene Set Enrichment Analysis

Gene set enrichment analysis (GSEA) was performed separately for each cell subtype using the clusterProfiler R package (v4.14.6) [27]. Genes from the pseudobulk DESeq2 analysis were ranked according to the Wald test statistic, with positive values indicating increased expression in HET relative to WILD. Gene sets were obtained from MSigDB using the msigdbr package for *Mus musculus* and included the Hallmark, Reactome, and Gene Ontology Biological Process (GO) collections. GSEA was performed using gene sets containing between 15 and 500 genes, and statistical significance was defined as a Benjamini–Hochberg-adjusted *p*-value < 0.05.

### Cell-cell communication analysis

Cell–cell communication differences between genotypes were analysed using CellChat v2.2.0 [28]. Raw UMI counts were normalized using CellChat’s internal procedure. Signalling inference was performed with CellChatDB.mouse, including protein, metabolic, and synaptic ligand–receptor categories. Signalling-relevant genes were selected using subsetData(), overexpressed genes and ligand–receptor pairs were identified, and communication probabilities were computed using the truncated mean method (trim = 0.1). Interactions involving cell populations represented by fewer than five nuclei or detected in fewer than two biological replicates were excluded. Pathway-level communication probabilities and aggregated signalling networks were subsequently calculated.

High-confidence ligand–receptor interactions (*p* < 0.05, communication probability > 0.05, ≥5% expressing cells) and signalling pathways (*p* < 0.05, communication probability > 0.02) were retained. HET and WILD communication networks were compared at both ligand–receptor and pathway levels. For each interaction, the difference in communication probability (HET − WILD) was calculated. Interactions detected only in HET or WILD were classified as gained or lost, respectively, interactions with an absolute difference in communication probability >0.05 were classified as changed, and those below this threshold were considered unchanged.

## Results

### Single-nucleus RNA-seq profiling of the CDKL5-deficient mouse hippocampus

To profile cell-subtype-specific transcriptional changes following CDKL5 deficiency, we performed droplet-based snRNA-seq on hippocampal tissue from adult female mice (Fig. 1a) [29,30]. Four mice were sequenced in total, two wild type (WT) and two heterozygous (HET), with a target of 3,000 nuclei to be sequenced per sample. Gel electrophoresis confirmed the genotype of the mice (Supplementary Fig. S1). After QC, we obtained 11,791 high-quality nuclei, with an average of 3,052 genes detected per cell. UMAP visualisation showed overlap between samples, with no obvious sample-specific separation (Fig. 1b).

**Figure 1.**
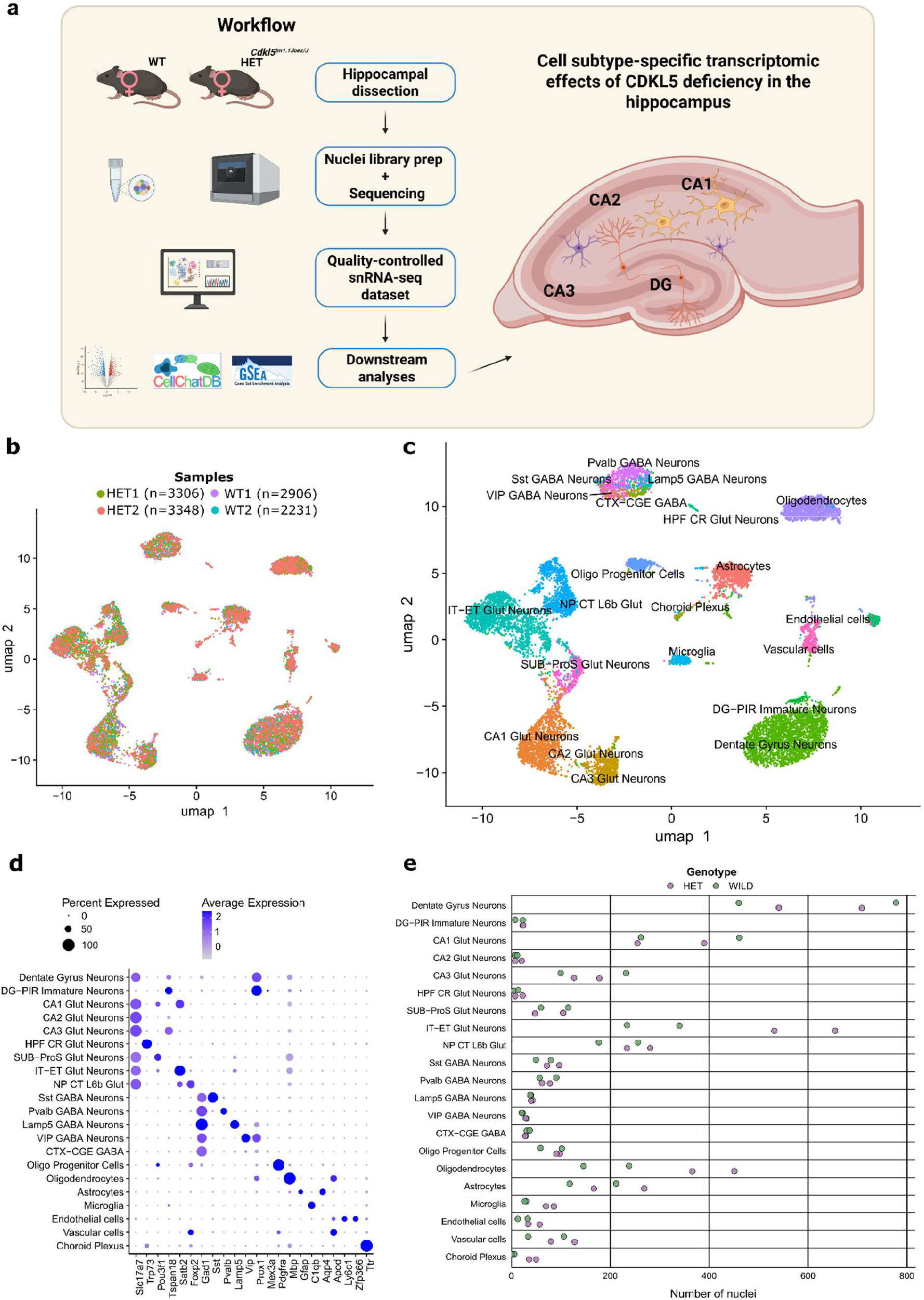
Single-nucleus transcriptomic profiling of the adult hippocampus in a heterozygous CDKL5-deficient mouse model. a) Schematic overview of the experimental and computational workflow used for single-nucleus RNA sequencing (snRNA-seq) of hippocampal tissue from wild-type (WT) and heterozygous (HET) mice. (b) Uniform Manifold Approximation and Projection (UMAP) embedding of all nuclei in the dataset, coloured by mouse of origin. c) UMAP embedding of all nuclei coloured and labelled according to annotated cell subtype. d) Dot plot showing the expression of canonical marker genes across annotated cell subtypes. Dot size represents the percentage of nuclei expressing each gene and colour intensity represents average expression. e) Number of nuclei assigned to each annotated cell subtype across WT and HET samples. Each point represents an individual sample.

We used MapMyCells [24] for reference-based annotation of our dataset by transferring cell labels from the Allen Whole Mouse Brain Atlas. Cell annotation confirmed the presence of canonical hippocampal cell subtypes, with an initial 10 broad cell subtypes identified in our dataset. These were further subdivided according to the Allen Brain Atlas annotation, resulting in 21 cell subtypes and a detailed representation of the cellular heterogeneity of the mouse hippocampus (Fig. 1c). We confirmed that hippocampal cell subtypes were consistently represented across samples (Supplementary Fig. S2), and examined region-specific marker expression to confirm the accuracy of cell annotations (Fig. 1d, Supplementary Fig. S3). Cell subtypes represented by fewer than 5 nuclei and that could not be integrated with other subtypes were discarded from downstream analyses.

Excitatory neurons represented approximately 65% of all cells and comprised several glutamatergic populations. Intratelencephalic–extratelencephalic-like Glut (IT–ET Glut) neurons were further resolved into subiculum-prosubiculum glutamatergic (SUB-ProS Glut) neurons, CA1 Glut neurons, CA2 Glut neurons, CA3 Glut neurons and near-projecting cortex layer 6b glutamatergic (NP CT L6b Glut) neurons. Nuclei that could not be assigned to one of these more specific populations were retained under the broader IT–ET Glut neuron label. Dentate gyrus (DG) neurons, dentate gyrus–piriform (DG-PIR) immature neurons, and hippocampal formation Cajal–Retzius glutamatergic (HPF CR Glut) neurons were also identified.

Inhibitory neurons accounted for approximately 8% of cells, and included parvalbumin (Pvalb), somatostatin (Sst), vasoactive intestinal peptide (Vip), and lysosome-associated membrane protein 5 (Lamp5) GABA neurons. Nuclei that could not be confidently assigned to a more specific subclass were retained under the broader Cortex - Caudal Ganglionic Eminence (CTX-CGE) GABA annotation. Glial populations represented approximately 21% of cells and comprised astrocytes, microglia, oligodendrocyte progenitor cells (OPCs) and oligodendrocytes. Additional non-neuronal populations included vascular cells, endothelial cells and choroid plexus cells. Following completed annotation, the most abundant subtypes in the dataset were DG neurons (2,488 nuclei), IT–ET Glut neurons (1,761), CA1 Glut neurons (1,368), oligodendrocytes (1,201) and NP CT L6b Glut neurons (948).

We next assessed whether the relative representation of cell subtypes differed between HET and WT mice. Most cell subtypes showed broadly similar numbers of nuclei between genotypes, although differences were observed for IT–ET Glut neurons, microglia and oligodendrocytes (Fig. 1e). Choroid plexus cells were also underrepresented in WT samples, which could be due to differences in the handling of the extracted hippocampus [31]. A slightly higher number of nuclei was recovered from HET samples, which may have influenced the observed differences in cell abundance. Thus, these trends should be interpreted with caution and further investigated in future studies with larger sample sizes.

Overall, our snRNA-seq dataset captures the major hippocampal cell populations and is appropriate for investigating the hippocampal gene expression changes associated with CDD.

### CDKL5 deficiency induces distinct transcriptional changes across hippocampal cell populations

To identify cell subtype-specific transcriptomic changes associated with CDKL5-deficiency, we performed differential expression analysis across conditions. A pseudobulk approach was used to account for inter-sample variability [32] using DESeq2’s pipeline, and results were filtered by FDR correction (adjusted *p* < 0.05), resulting in a total of 75 significant differentially expressed genes (DEGs) across subtypes (Fig. 2a, Supplementary Table S1). PCA of cell type-specific pseudobulked expression profiles identified several cell subtypes where HET samples grouped closer to each other than to WT samples (e.g. IT-ET Glut neurons, oligodendrocytes, microglia), suggesting that genotype may contribute substantially to transcriptional variation in these populations (Supplementary Figs. S4-5). Differential expression analysis identified a higher number of DEGs in IT-ET Glut Neurons (38) and Dentate Gyrus Neurons (22).

**Fig. 2.**
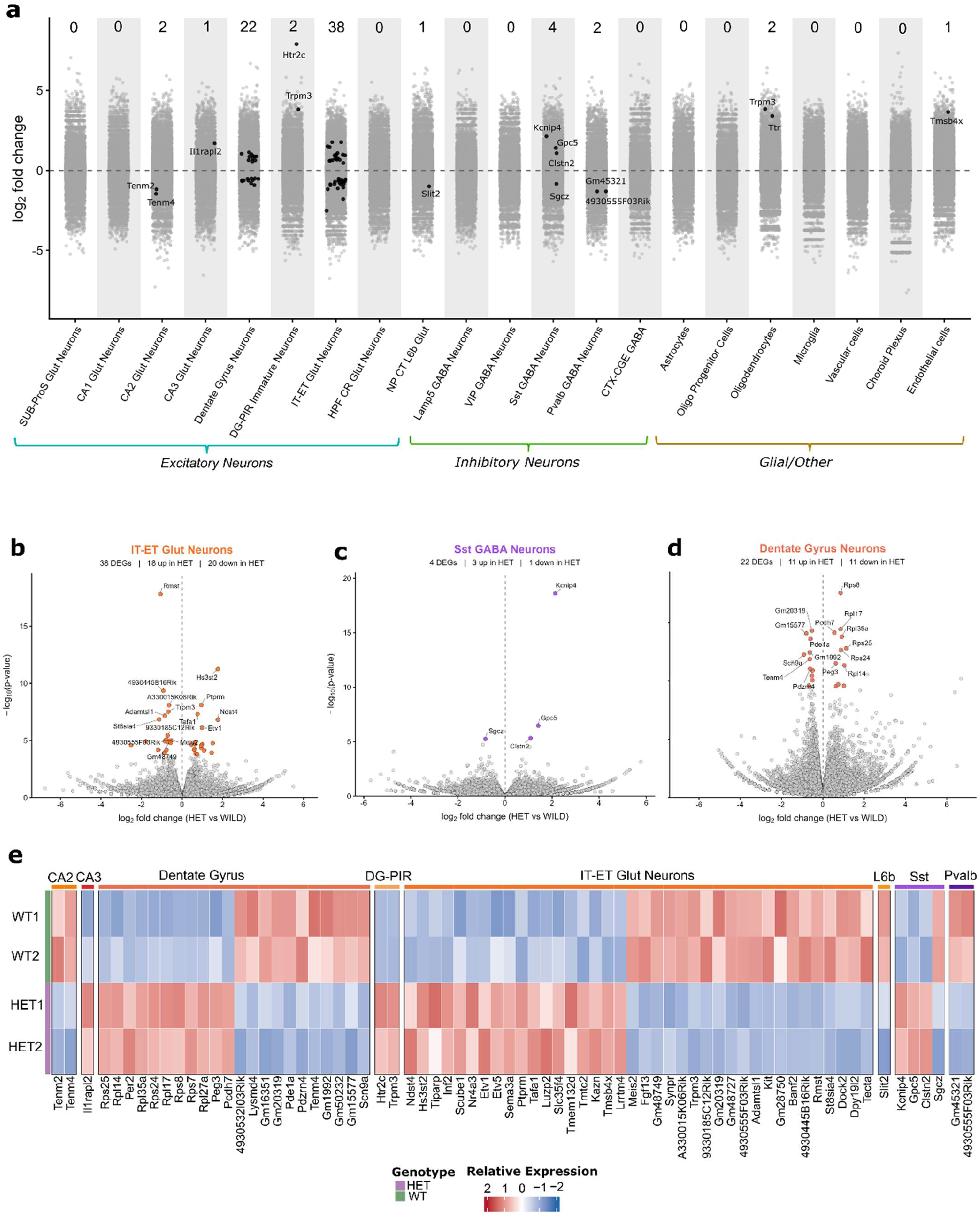
Cell subtype-specific differential gene expression in the CDKL5-deficient hippocampus. a) Summary of DESeq2 pseudobulk differential expression analysis across annotated hippocampal cell subtypes. Each point represents a gene plotted by log_2_ fold change for the comparison between HET and WT samples, with positive values indicating higher expression in HET. Significantly differentially expressed genes are highlighted in black (p_adj<0.05), and the number above each subtype indicates the total number of significant DEGs identified. Cell subtypes are grouped into excitatory neurons, inhibitory neurons and glial/other populations. (b-d) Volcano plots showing differential gene expression in (b) IT–ET Glut neurons, (c) Sst GABA neurons and (d) Dentate Gyrus neurons. Labelled genes indicate selected significant DEGs (p_adj<0.05). (e) Heatmap showing the relative expression of selected differentially expressed genes across hippocampal cell subtypes and samples. Rows represent individual samples and columns represent genes. Colours indicate Z-score normalised expression within each subtype-pseudobulk profile.

IT-ET Glut Neurons (Fig. 2b) displayed downregulation of genes linked to synaptic function and signalling(e.g., *Synpr, Trpm3,* and *Fgf13*), with upregulation of genes involved in axon guidance, cell adhesion, and synaptic organisation (e.g., *Sema3a*, *Lrrtm4*, and *Ptprm* [33–35]), suggesting transcriptional changes affecting neuronal connectivity and synaptic function.. We also observed upregulation of transcription factors and genes associated with activity-dependent programs (e.g., *Etv1/Etv5* and *Nr4a3* [36,37]), and downregulation of *Rmst,* a long non-coding RNA involved in neurogenesis and neuronal differentiation [38].

DG neurons displayed a prominent ribosomal transcriptional signature (Fig. 2d). Eight ribosomal transcripts (*Rps7*, *Rps8*, *Rps24*, *Rps25*, *Rpl14*, *Rpl17*, *Rpl27a*, and *Rpl35a*) were upregulated in HET mice, suggesting altered ribosomal and translation-related processes. Additional transcriptional changes in DG neurons again involved genes associated with neuronal signalling, cell adhesion, and synaptic organisation, including upregulation of *Pcdh7* and downregulation of *Pde1a*, *Tenm4*, and *Pdzrn4*. Notably, reduced *Tenm4* expression was also observed in CA2 Glut neurons, together with downregulation of the related teneurin gene *Tenm2* (Fig. 2a). Together, these findings indicate that CDKL5 deficiency is associated with changes in both ribosomal processes and neuronal connectivity-related genes within hippocampal excitatory populations.

Sst GABA neurons were the most affected inhibitory neuronal subtype (Fig. 2c), with upregulation of *Kcnip4, Gpc5,* and *Clstn2*, genes associated with ion-channel regulation [39], neuron-neuron signalling [40,41], and synaptic organisation [42], and downregulation of *Sgcz,* associated with postsynaptic organisation [43]. Together, these changes suggest that CDKL5 deficiency may also alter synaptic signalling and organisation within Sst GABA neurons.

Overall, we identified more DEGS in neuronal populations (Fig. 2e) compared to non-neuronal cells. Oligodendrocytes displayed upregulation of *Ttr* and *Trpm3* (Fig. 2a), while endothelial cells showed an upregulation of *Tmsb4x* (Fig. 2a), a gene associated with brain microvascular endothelial function [44].

As a complementary resource, we also performed single-nucleus differential expression analysis using MAST to provide a broader overview of transcriptional alterations across individual cell populations [25] (Supplementary Figs. S6-7, Supplementary Table S2). These results are provided as a supplementary resource. Statistical inference and biological interpretation throughout the paper were based on pseudobulk DESeq2 analysis, which accounts for biological replicates and reduces the risk of false-positive findings [32].

### Metabolic, inflammatory and stress-response programmes are enriched in the CDKL5-deficient hippocampus

To investigate whether biological processes were altered in CDKL5-deficient mice beyond changes detectable at the level of individual genes, we performed gene set enrichment analysis (GSEA) using ranked pseudobulk differential expression statistics. This approach allowed us to identify coordinated pathway-level changes across the full ranked expression profile, including changes that may not be captured by analyses based only on significantly differentially expressed genes.

Hallmark analysis [45] identified 149 significant cell subtype–pathway enrichments (adjusted *p* < 0.05), with a strong predominance (144/149) of positive enrichment in HET mice (Fig. 3a, Supplementary Table S3). The highest number of pathway-level responses were observed in neuronal populations, particularly NP CT L6b and CA1 Glut neurons, although several non-neuronal populations, including endothelial cells, vascular cells and oligodendrocytes, also showed multiple significantly enriched pathways.

**Fig. 3.**
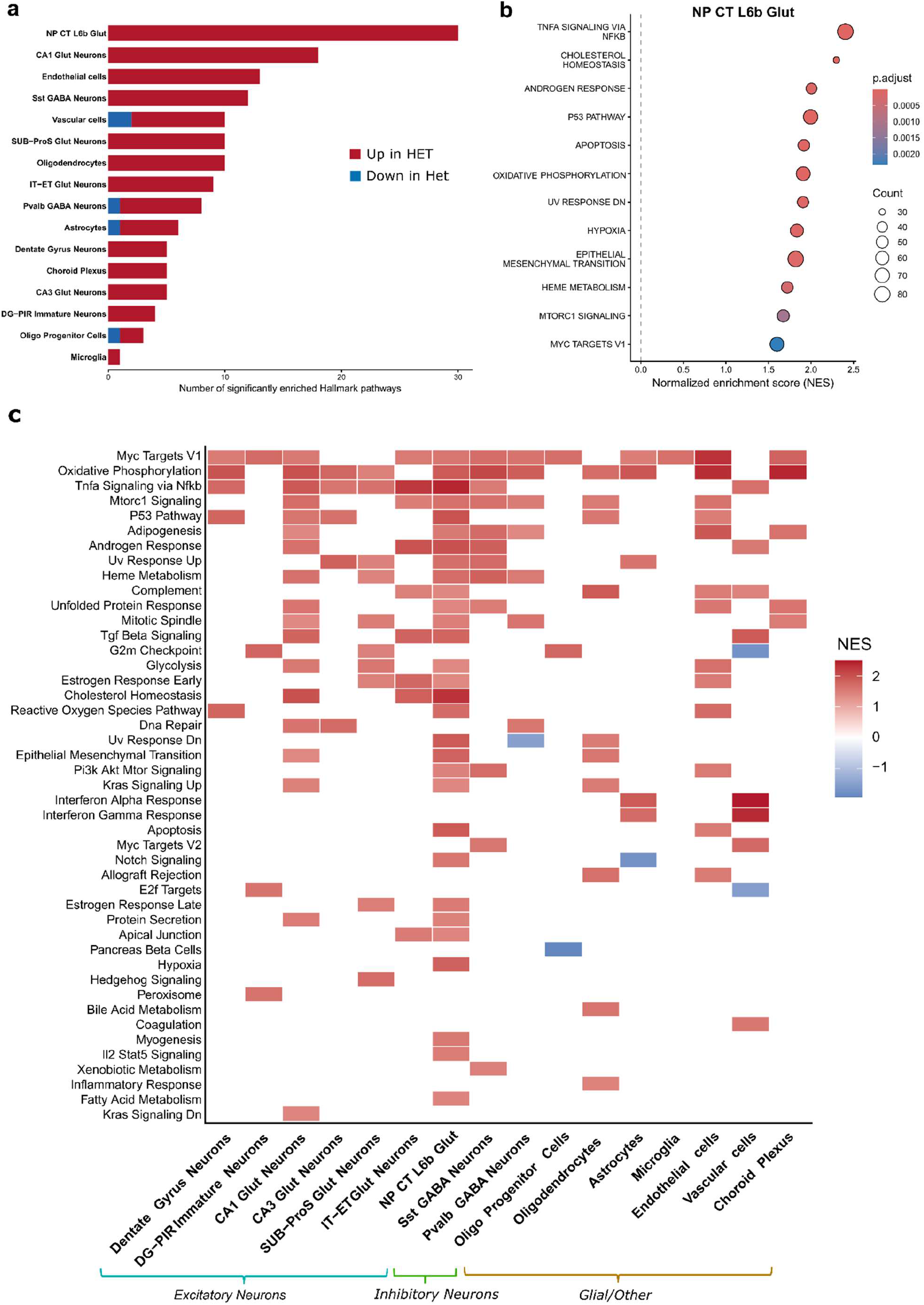
Cell subtype-specific enriched Hallmark pathways in the CDKL5-deficient hippocampus. (a) Number of significantly enriched Hallmark pathways (p_adj<0.05) across hippocampal cell subtypes. Pathways with positive normalized enrichment scores (NES) are shown as up in HET (orange), while pathways with negative NES are shown as up in WT (blue). (b) Dot plot showing significantly enriched Hallmark pathways in NP CT L6b Glut neurons. Dot size represents the number of genes contributing to each gene set and colour indicates the adjusted p-value. (c) Heatmap showing significantly enriched Hallmark pathways across hippocampal cell subtypes. Colours indicate the NES, with positive values representing enrichment in HET and negative values representing enrichment in WT. Cell subtype labels are grouped according to cell class (Excitatory Neurons, Inhibitory Neurons, or non-neuronal (Glial/Other)).

NP CT L6b Glut neurons presented the strongest Hallmark response (Fig. 3b), with enrichment of inflammatory responses (TNFα signalling via NF-κB) alongside hypoxia and apoptosis-related programmes, reflecting cellular stress. Metabolic alterations included enrichment of oxidative phosphorylation and cholesterol homeostasis, while mTORC1 signalling, p53 signalling and MYC targets suggested additional changes in cellular growth and transcriptional regulation.

Similar molecular programmes were enriched across multiple cell populations (Fig. 3c). Oxidative phosphorylation and TNFα signalling via NF-κB were frequently enriched across subtypes, suggesting widespread changes in cellular metabolism and inflammatory responses. MYC targets and mTORC1 signalling were also enriched across several populations, pointing to changes in cellular growth and transcriptional regulation. In addition, stress-response pathways, including p53 signalling, the unfolded protein response and reactive oxygen species pathways, were altered across multiple cell subtypes.

Together, these findings indicate that CDKL5 deficiency could be associated with widespread alterations in metabolic, inflammatory, stress-response and growth-related programmes across hippocampal cell types.

### Cell subtype-specific enrichment of neuronal development and synaptic processes in CDKL5 deficiency

While Hallmark analysis provided an overview of broad molecular programmes altered in CDKL5 deficiency, these gene sets represent relatively general cellular processes. We therefore next performed GSEA using Gene Ontology Biological Process (GO-BP) [45,46] gene sets to investigate the affected biological processes at a more detailed and cell subtype-specific level.

GO-BP analysis revealed a strong cell subtype-specific response to CDKL5 deficiency, with distinct biological processes altered across individual populations (Fig. 4a). NP CT L6b Glut neurons displayed the largest number of significantly enriched GO-BP terms, followed by oligodendrocytes and CA1 Glut neurons. However, several related processes were also altered across multiple subtypes, suggesting that some responses to CDKL5 deficiency were shared between hippocampal cell populations.

**Fig. 4.**
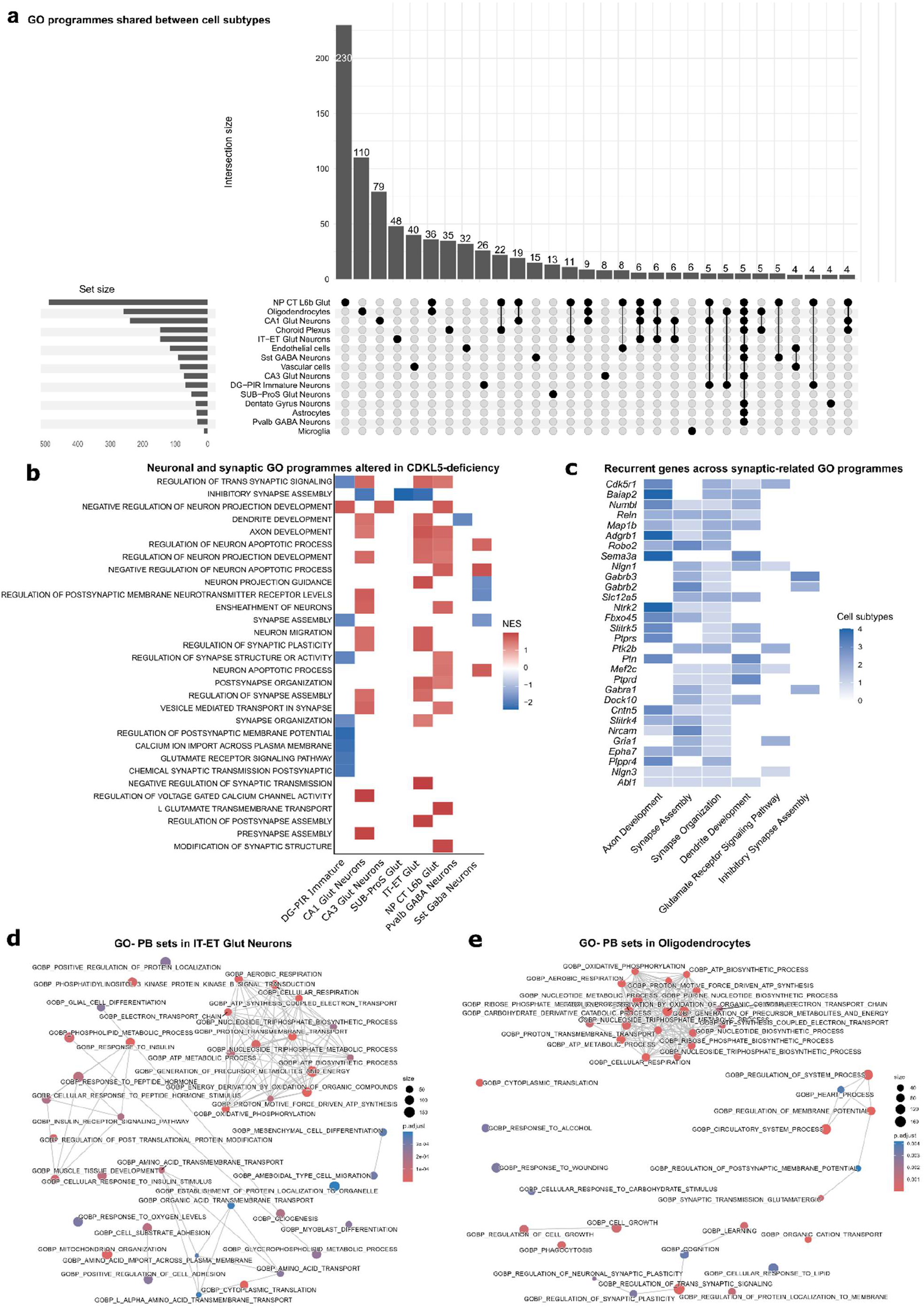
Enriched Gene Ontology biological processes across hippocampal cell subtypes in CDKL5 deficiency. (a) UpSet plot showing shared significantly enriched Gene Ontology Biological Processes (GO-BP) across hippocampal cell subtypes. Bars indicate the number of GO-BPs shared between the indicated cell subtypes, while horizontal bars show the total number of significantly enriched GOBPs within each subtype. (b) Heatmap showing selected neuronal and synaptic GO-BPs across neuronal subtypes. Colours indicate the normalized enrichment score (NES), with positive values representing enrichment in HET and negative values representing enrichment in WT. (c) Heatmap showing recurrent leading-edge genes across selected synaptic GO-BPs. Colour intensity indicates the number of cell subtypes in which each gene contributed to the leading edge of the indicated pathway. (d-e) Enrichment map showing significantly enriched GO-BPs in (d) IT-ET Glut neurons and (e) oligodendrocytes. Node size represents the number of genes within each GO-BP and colour indicates the adjusted p-value.

Given the predominantly neuronal expression and function of CDKL5 [6], we focused on GO-BP terms related to neuronal function within neuronal subtypes (Fig. 4b). Alterations in axon and dendrite development, neuron projection organisation and synaptic function were observed across several neuronal populations, but their enrichment patterns differed between subtypes. DG-PIR Immature Neurons showed negative enrichment of several synaptic processes, including trans-synaptic signalling, synapse organisation and glutamatergic postsynaptic signalling. In contrast, related neuronal and synaptic processes showed positive enrichment in excitatory populations including CA1 Glut, IT-ET Glut and NP CT L6b Glut neurons. Changes in inhibitory neurons were more restricted, with Pvalb and Sst GABA neurons showing alterations in processes related to dendrite development, neuron projection guidance, synapse organisation and postsynaptic receptor regulation (Fig. 4b). Together, these findings suggest that CDKL5 deficiency affects overlapping aspects of neuronal development and synaptic function, but that the transcriptional response differs substantially between neuronal subtypes.

To identify the genes that contribute to these altered neuronal processes, we examined the leading-edge genes underlying these selected synaptic and neuronal GO-BP enrichments (Fig. 4c). Several genes were repeatedly present across multiple related processes and cell subtypes, including genes involved in neurite development and guidance (*Sema3a, Robo2, Map1b*), synaptic organisation (*Baiap2, Nlgn1, Nlgn3*), neurotransmitter receptor function (*Gabrb2, Gabrb3, Gabra1*), and neuronal signalling (*Ntrk2, Cdk5r1, Slc12a5*). This indicates that the enrichment of GO-BP terms related to neuronal and synaptic function is partly driven by recurrent genes across multiple cell types.

Finally, enrichment-map analysis was used to examine the broader relationships between significantly altered GO-BP terms (Fig. 4d-e, Supplementary Table 4). In IT-ET Glut neurons, enriched processes formed clusters associated with mitochondrial respiration and ATP synthesis, insulin and peptide hormone responses, and amino acid transport (Fig. 4d). Oligodendrocytes showed prominent enrichment of metabolic processes, particularly mitochondrial respiration and ATP synthesis, alongside smaller clusters of GO-BP terms related to synaptic signalling and plasticity (Fig. 4e).

Overall, GO-BP analysis identified neuronal development and synaptic organisation as recurrent transcriptional signatures of CDKL5 deficiency across hippocampal neuronal populations. The identification of shared leading-edge genes further highlights common transcriptional changes associated with neuronal connectivity and synaptic function, while the prominent enrichment of metabolic pathways in oligodendrocytes also suggests a glial response to an altered neuronal environment.

### Widespread alterations in inferred cell–cell communication in the CDKL5-deficient hippocampus

To further investigate the molecular alterations associated with CDKL5 deficiency, we next examined predicted cell–cell communication between hippocampal cell populations using CellChat [28]. This analysis revealed widespread differences in both the number and strength of inferred interactions between HET and WT samples, while the magnitude and direction of these changes varied considerably between sender and receiver populations (Fig. 5a). Changes in interaction number were particularly prominent among neuronal populations, while differences in overall interaction strength showed a more heterogeneous pattern, suggesting that CDKL5 deficiency affects both the extent and relative strength of intercellular signalling in a cell subtype-dependent manner.

**Fig. 5.**
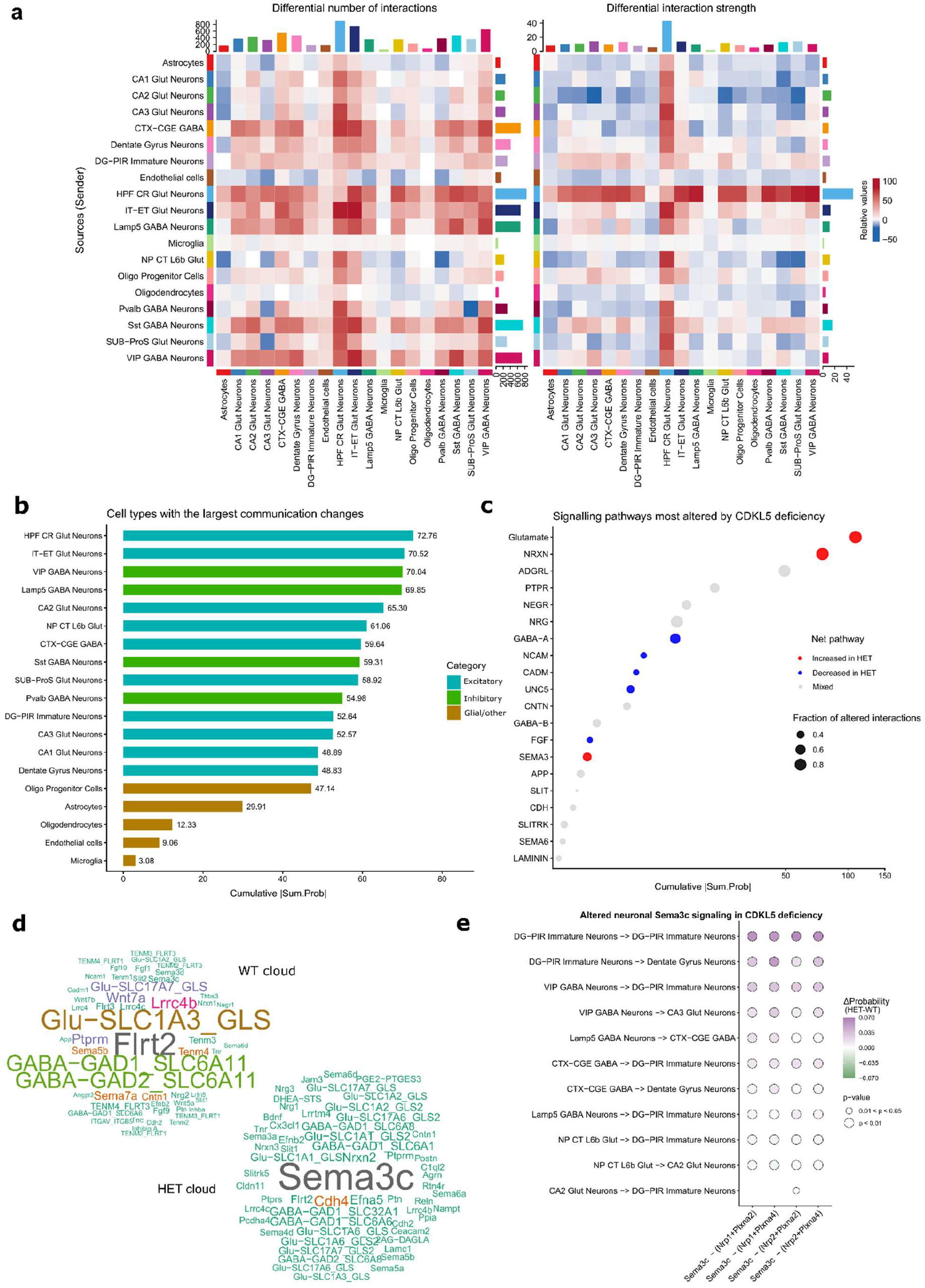
CellChat predicts altered cell-cell communication in the CDKL5-deficient hippocampus. (a) Heatmaps of inferred changes in the number of cell-cell interactions (left) and interaction strength (right). Cell subtypes in the x-axis are receivers of the information flow (expressing receptors), while those in the y-axis are senders (expressing ligands). Positive relative values (red shading) indicate increased communication in HET mice, while negative values (blue shading) indicate decreased communication. (b) Bar plot ranking cell subtypes by the summed absolute change in significant communication probabilities (Σ|Δprob|) across all signalling pathways. (c) Dot plot ranking signalling pathways by the cumulative absolute change in communication probability across all cell subtypes. Dot size represents the fraction of altered interactions within each pathway, while colour indicates whether signalling was predominantly increased in HET (red), decreased in HET (blue), or showed mixed directional changes (grey). d) Word clouds showing enriched signalling terms among differential ligand–receptor interactions in WT and HET mice. Text size represents the relative enrichment score of each ligand. (e) Dot plot showing altered neuronal Sema3c signalling between selected cell subtypes. Positive Δprob values indicate increased signalling in HET mice, while negative values indicate decreased signalling. Dot colour represents the change in communication probability between conditions and dot size indicates statistical significance (all dots have p<0.05).

To identify the populations contributing most strongly to these changes, we ranked cell subtypes according to the total absolute difference in inferred communication probability (Fig. 5b). HPF CR Glut neurons showed the largest overall change, followed by IT–ET Glut, Vip GABA and Lamp5 GABA neurons, while most non-neuronal populations showed comparatively smaller differences. This indicates that the major alterations in predicted cell–cell communication were concentrated within neuronal populations.

At the pathway level, glutamatergic and NRXN (neurexin) signalling showed some of the largest communication changes, with a general increase in HET mice (Fig. 5c). ADGRL (latrophilin) signalling also showed substantial alterations, although these involved a mixture of increased and decreased interactions in the communicating cell subtypes. In contrast, GABA-A signalling and several pathways associated with cell adhesion and neuronal connectivity, including NCAM (Neural Cell Adhesion Molecule), CADM (Cell Adhesion Molecules), UNC5 and FGF (Fibroblast Growth Factor) signalling, showed an overall reduction in HET mice.

Analysis at the ligand–receptor level further revealed distinct signalling patterns between conditions (Fig. 5d). WT samples showed greater enrichment of glutamate and FLRT interactions, whereas HET samples were characterised by prominent *Sema3c* signalling alongside changes in several cell adhesion and synaptic-associated interaction

Given the prominence of SEMA3 signalling in HET mice, we next examined neuronal *Sema3c* interactions in greater detail (Fig. 5e). These interactions were primarily mediated through *Nrp1* or *Nrp2* together with *Plxna2* or *Plxna4*, and showed marked sender–receiver specificity. DG-PIR Immature Neurons showed some of the strongest increases as signalling sources, with enhanced predicted communication towards DG-PIR Immature Neurons, DG neurons and other neuronal populations. Increased SEMA3 signalling was also observed from Vip GABA, CTX-CGE GABA and Lamp5 GABA neurons, whereas a smaller number of interactions were stronger in WT mice. These findings suggest that altered SEMA3 signalling in CDKL5 deficiency is concentrated within specific neuronal communication networks rather than representing a uniform increase across the pathway

Overall, CellChat analysis identified extensive cell subtype- and pathway-specific alterations in predicted intercellular communication, including increased glutamatergic and NRXN signalling, reduced GABA-A and adhesion-associated signalling, and selective increases in Sema3c-mediated communication involving DG-PIR Immature Neurons and other neuronal populations. These findings suggest that CDKL5 deficiency may alter the balance of excitatory, inhibitory and guidance-related signalling within hippocampal neuronal networks.

## Discussion

In this study, we present the first single-nucleus transcriptional atlas of the hippocampus from heterozygous female mice carrying a deletion of exon 6 of *Cdkl5*, a mouse model of CDD. We identify widespread changes in inferred cell-cell communication, involving neuronal and glial populations, together with potential transcriptional alterations associated with synaptic structure and organisation. Together, the generated data strengthen our understanding of the cell subtype-specific transcriptional alterations associated with CDKL5 deficiency in the murine hippocampus, providing a resource for potential biomarker and therapeutic target discovery in CDD.

Previous studies have investigated gene expression changes associated with CDD, but these have largely relied on bulk transcriptomic approaches, which mask the cellular heterogeneity of the brain [10,12,47]. Single-cell omics technologies have therefore enabled high-resolution characterisation of gene expression across individual cell subtypes [20,48]. These technologies have been previously used to assess transcriptomic changes in related neurodevelopmental and epilepsy murine models, including Rett syndrome, fragile X syndrome and temporal lobe epilepsy [15,49,13].

Mouse models of CDD recapitulate several features of the human disorder and have previously demonstrated distinct contributions of neuronal populations to CDD-associated phenotypes [18,50]. The present study used heterozygous female mice, reflecting the genetic context observed in most individuals with CDD [2], and extends previous molecular characterisation of these models by examining transcriptional alterations across individual hippocampal cell subtypes.

CDKL5 is predominantly expressed in excitatory and inhibitory neurons in the mouse brain, with comparatively low expression in non-neuronal populations [6], and its loss alters excitatory-inhibitory balance. Consistent with this, neuronal subtypes in our dataset displayed the greatest number of differentially expressed genes across conditions, with IT–ET Glut and DG neurons showing the strongest transcriptional alterations. Several of the differentially expressed genes have established roles in neuronal maturation, morphology and synaptic organisation. For example, reduced expression of FGF13, which regulates axonal development and voltage-gated sodium channels, may be relevant to neuronal morphology and excitability, particularly given pathogenic variants in FGF13 have been associated with developmental and epileptic encephalopathy [51,52].

Conversely, several genes with increased expression in IT–ET Glut neurons are associated with neuronal structural remodelling, such as *Sema3a,* an axon-guidance molecule that regulates neuronal polarity [33], *Etv1/Etv5,* ETS-family transcription factors that participate in activity-dependent transcriptional programmes associated with neuronal maturation [37] and hippocampal dendritic arborisation and spine development [53], respectively; and *Lrrtm4,* a postsynaptic adhesion molecule involved in excitatory synapse formation and function [34]. Together, these transcriptional alterations suggest that CDD may induce coordinated changes in pathways controlling neuronal morphology, maturation and synaptic organisation within IT–ET Glut neurons.

DG neurons also showed increased expression of multiple ribosomal protein genes in HET mice, potentially reflecting a cellular response to increased translational demands. This observation is particularly relevant given previous evidence linking CDKL5 deficiency with disruption of mTOR– ribosomal signalling. CDKL5-deficient mice have previously shown altered AKT/ribosomal protein S6 signalling and reduced phosphorylation of ribosomal protein S6 [9], while cell subtype-specific deletion of CDKL5 has been shown to alter components of the mTOR pathway, including the downstream S6 kinase p70S6K1 [54]. The apparent coordinated increase in ribosomal transcripts observed may therefore reflect cell subtype-specific dysregulation or compensatory remodelling of translational pathways within the DG, although direct assessment of protein synthesis would be required to establish its functional consequence.

GSEA using Hallmark and GO-BP sets revealed recurrent transcriptional changes associated with cellular metabolism, growth regulation, and stress responses across hippocampal cell subtypes. Among these, metabolic programmes were particularly prominent, with oxidative phosphorylation showing widespread enrichment. This is particularly interesting given previous evidence of altered mitochondrial function in CDD [47]. CDKL5-deficient mouse brains have shown reduced ATP levels and altered mitochondrial electron transport chain function [55], while patient-derived CDKL5-deficient neuronal cells display abnormalities in mitochondrial respiratory-chain activity and bioenergetics [47,56]. These findings could suggest that disruption of cellular energy-associated programmes may represent a recurrent molecular feature of CDKL5 deficiency, although the transcriptional enrichment observed here does not necessarily imply measurement of increased mitochondrial activity and future studies are needed to determine the functional significance of these changes.

Across neuronal subtypes, recurrent alterations were observed in pathways associated with neuronal projection development, axonal and dendritic organisation, synapse assembly, synaptic signalling and neurotransmission. However, the specific pathways and their direction of enrichment differed between subtypes, indicating that CDKL5 deficiency does not produce a uniform neuronal response. Instead, related aspects of neuronal structure and synaptic function appear to be affected through distinct transcriptional programmes across hippocampal populations. Similar transcriptomic alterations have also been reported in *Cdkl5*-mutant zebrafish, where bulk RNA-seq identified changes in gene expression associated with pathways related to neuronal development, cytoskeletal organisation, extracellular-matrix interactions and synaptic signalling [12]. Together, these findings support disruption of neuronal development and connectivity-related molecular programmes as a recurrent feature of CDKL5 deficiency, while our data further resolve these alterations to specific hippocampal cell subtypes.

Despite having comparatively few significantly differentially expressed genes, Sst and Pvalb GABA neurons showed enrichment of GO-BP terms related to dendritic development, synapse organisation and postsynaptic receptor regulation, suggesting that coordinated transcriptional changes may be detected at the pathway level even when few individual genes reach conventional differential-expression thresholds [57]. This may be particularly relevant given previous studies demonstrating that both excitatory and inhibitory neuronal populations contribute to CDD-associated phenotypes [7,58]. Similarly, single-nucleus transcriptomic analysis of Rett syndrome, a related X-linked neurodevelopmental disorder, identified neuronal and synaptic pathway alterations across both excitatory and inhibitory populations [16].

Despite comparatively low CDKL5 expression in non-neuronal cells [6], oligodendrocytes exhibited a prominent GO-BP enrichment profile, potentially consistent with a glial response associated with CDKL5 deficiency. Among the significant processes detected, the dominant clusters comprised gene sets related to oxidative phosphorylation, aerobic and cellular respiration, the electron transport chain, ATP biosynthesis and metabolism, and altered cytoplasmic translation. This metabolic signature is particularly relevant to oligodendrocyte biology, as these cells not only generate and maintain myelin but also provide essential metabolic support to the axons they bind, and oligodendrocyte energy metabolism can influence both myelin maintenance and axonal support [59,60]. The enrichment of metabolic programmes in the oligodendrocytes of *Cdkl5*-deficient mice may therefore indicate altered energetic demands or an altered neuron–glia metabolic environment.

Cell-cell communication within the hippocampus also appeared widely altered in our dataset, with neuronal subtypes dominating these changes. Alterations were not restricted to a single neuronal population, but involved multiple excitatory and inhibitory subtypes, suggesting widespread remodelling of predicted intercellular signalling across the CDKL5-deficient hippocampus. Among the altered pathways, predicted SEMA3C signalling was particularly prominent in HET samples, with further analysis identifying DG-PIR immature neurons, DG neurons and inhibitory populations including Vip and Lamp5 interneurons as major contributors to this signal. Class 3 semaphorins (SEMA3s) are secreted guidance molecules with important roles in neuronal development, axonal and dendritic growth and the establishment of neuronal connectivity [61]. SEMA3C has previously been shown to regulate axonal growth within the hippocampal formation, including the development of septohippocampal projections [62]. Interestingly, our differential expression analysis also identified increased *Sema3a* expression in IT–ET Glut neurons, suggesting that altered semaphorin-associated signalling is not restricted to the SEMA3C interactions predicted by CellChat. SEMA3A regulates axonal and dendritic development in hippocampal neurons [33] and has also been implicated in dendritic branching and spine maturation, including regulation of PSD95 and synapsin organisation [63].

These findings may be particularly relevant in the context of previously described synaptic and dendritic abnormalities in CDD. *Cdkl5*-deficient mice show reduced dendritic complexity within hippocampal pyramidal neurons [30], while loss of CDKL5 has also been associated with reduced spine stability, decreased PSD95-positive synaptic puncta and impaired synaptic plasticity [64]. The convergence of increased *Sema3a* expression with increased predicted SEMA3C signalling may therefore indicate broader dysregulation of semaphorin-mediated pathways involved in neuronal connectivity and synaptic organisation in the CDKL5-deficient hippocampus. However, whether these changes contribute directly to the synaptic phenotype of CDD or represent compensatory responses to altered neuronal connectivity remains to be determined.

Teneurins are brain-enriched adhesion molecules with important roles in neuronal connectivity and synapse formation, through interactions with their latrophilin receptors [65,66]. Alterations in teneurin signalling have been reported in neurological disorders including the epilepsies [67–69], and seizure activity has been shown to modify expression of teneurins [13,67]. This makes the reduced expression of *Tenm2* and *Tenm4* in CA2 Glut neurons, together with reduced *Tenm4* in DG neurons, particularly interesting. These alterations may contribute to the abnormalities in dendritic growth and excitatory synapse organisation described in CDD [64,70], given the established roles of teneurins in neuronal development and circuit formation [71,72]. However, the functional direction of these changes remains unclear. Reduced teneurin expression could contribute to aberrant circuit formation and thereby increase susceptibility to network hyperexcitability, but it could alternatively represent a compensatory response limiting excitatory connectivity in an already hyperexcitable network.

This possibility is particularly relevant given the differences between the seizure phenotype of individuals with CDD and commonly used mouse models. Whereas early-onset, often treatment-resistant epilepsy is a defining clinical feature of CDD [2], *Cdkl5*-deficient mice do not consistently reproduce this early spontaneous seizure phenotype [18,73]. Nevertheless, aged heterozygous female mice can develop spontaneous seizure-like events and epileptic spasms, with seizure prevalence increasing with age [73]. More recent studies using the same *Cdkl5* exon 6 deletion model have also demonstrated increased seizure susceptibility, including susceptibility to audiogenic seizures [74]. This suggests that *Cdkl5*-deficient networks may possess an underlying susceptibility to hyperexcitability even when spontaneous seizures are absent at earlier ages. Whether the altered teneurin expression identified here contributes to this susceptibility or instead represents a protective adaptation that limits seizure emergence remains unknown and will require functional investigation to assess their contribution to the phenotype, if any.

Limitations of the present study should also be acknowledged. Single-nucleus RNA sequencing captures nuclear transcripts and therefore provides a limited representation of cytoplasmic mRNA compared with whole-cell approaches. The relatively small number of biological replicates may also limit the detection of transcriptional changes in less abundant cell populations. In addition, approaches such as GSEA and CellChat provide inferred pathway enrichment and cell–cell communication patterns and therefore require functional validation. Increasing sample size and the number of nuclei profiled in future studies may help identify additional changes in low abundance cell subtypes. Our analysis was also restricted to the hippocampus, and profiling additional brain regions such as the cortex and amygdala may provide a broader view of the cellular consequences of CDKL5 deficiency. Integration with spatial transcriptomic or multimodal approaches could further help localise these transcriptional changes within hippocampal substructures and relate them to functional cellular phenotypes.

An important consideration is the use of heterozygous female mice in our study. As *CDKL5* is located on the X chromosome and undergoes X-chromosome inactivation, individual cell populations contain a mixture of cells expressing either functional or mutant CDKL5 [73]. The transcriptional changes identified here therefore represent the combined response of these two populations and do not allow us to determine whether individual signals arise predominantly from *Cdkl5*-deficient cells or from responses occurring in neighbouring cells retaining functional CDKL5. Future studies able to distinguish these populations at single-cell resolution will be important to separate direct cell-autonomous effects of CDKL5 loss from compensatory or non-cell-autonomous responses.

In conclusion, in the present study we profiled cell subtype-specific transcriptional changes in the hippocampus of a CDD mouse model. Our findings reveal distinct transcriptional responses across neuronal and glial populations, with recurrent alterations involving neuronal structure, ribosomal metabolism, synaptic organisation, cellular metabolism and predicted intercellular signalling. This dataset provides a valuable resource for understanding the cellular consequences of CDKL5 deficiency and establishes a foundation for future mechanistic studies, biomarker discovery and therapeutic investigation in CDD.

## Acknowledgments

The authors would like to thank Dr. Amaya Sanz Rodriguez and the FutureNeuro operations team for their valuable support throughout this work.

## Data Availability Statement

Raw FASTQ files and processed expression matrices are submitted to GEO and will be accessible upon publication. Supplementary files supporting the conclusions of this article will be included within the article and its supplementary materials. Code to reproduce the analyses will be available on GitHub.

## Ethics Statement

All procedures involving animals were performed in compliance with EU Directive 2010/63/EU on the protection of animals used for scientific purposes and procedures were approved by RCSI University of Medicine and Health Sciences’ Research Ethics Committee (REC 1587), and under license from the Ireland Health Products Regulatory Authority (AE19127/P087).

## Author Contributions

Original conception of the work: O.M. and D.C.H. Contribution and design of the experiment: O.M., J.V.S, S.E and A.S. Collection of data: J.V.S, S.E., A.S, and O.M. Analysis and interpretation: J.V.S, S.E,N.MC.C, D.C.H and O.M. Drafting the article: J.V.S, S.E, N.MC.C, D.C.H, and O.M. Revising the article: J.V.S, S.E, B.E.-M, J.H, A.S, M.H, D.C.H, and O.M. All authors approved the final version of the manuscript.

